# Pervasive SUMOylation of heterochromatin and piRNA pathway proteins

**DOI:** 10.1101/2022.08.15.504007

**Authors:** Maria Ninova, Hannah Holmes, Brett Lomenick, Katalin Fejes-Tóth, Alexei A. Aravin

## Abstract

Genome regulation involves complex and highly regulated protein interactions that are often mediated through post-translational modifications (PTMs). SUMOylation – the covalent attachment of the <u>s</u>mall <u>u</u>biquitin-like <u>mo</u>difier (SUMO) – is a conserved PTM in eukaryotes that has been implicated in a number of essential processes such as nuclear import, DNA damage repair, transcriptional control, and chromatin organization. In *Drosophila*, SUMO is essential for viability and its depletion from the female germline causes infertility associated with global loss of heterochromatin, and illicit upregulation of transposons and lineage-inappropriate genes. However, the specific targets of SUMO and its mechanistic role in different cellular pathways are still poorly understood. Here, we developed a proteomics-based strategy to characterize the SUMOylated proteome in *Drosophila* that allowed us to identify ~1500 SUMO sites in 843 proteins in the fly ovary. A high-confidence set of SUMOylated proteins is highly enriched in factors involved in heterochromatin regulation and several different aspects of the piRNA pathway that represses transposons, including piRNA biogenesis and function. Furthermore, we show that SUMOylation of several piRNA pathway proteins occurs in a Piwi-dependent manner, indicating a functional implication of this modification in the cellular response to transposon activity. Together, these data highlight the impact of SUMOylation on epigenetic regulation and reveal an unexpectedly broad role of the SUMO pathway in the cellular defense against genomic parasites. Finally, this work provides a valuable resource and a system that can be adapted to the study of SUMOylation in other *Drosophila* tissues.

## Introduction

Post-translational modifications (PTM) impact a broad range of cellular functions such as protein turnover, localization to different subcellular compartments, or specific interactions, and are utilized in the regulation of various molecular pathways. PTMs involve the covalent attachment of diverse moieties from small chemical groups to entire modifier proteins to the main polypeptide chain. The best-known protein modifier is ubiquitin – a ~9kDa unit that gets conjugated to target lysine side chains via an isopeptide bond between its C-terminal carboxyl group and the lysine epsilon amino group. Following the discovery of Ubiquitin, other small proteins that can act as modifiers have emerged, including the small ubiquitin-like modifier SUMO. SUMO is a ~11 kDa protein that shares structural and sequence homology with Ubiquitin and similarly gets covalently attached to target lysines. However, SUMO conjugation is mediated by different enzymes and serves distinct and non-redundant functions (reviewed in ref.^1^). In brief, the SUMOylation cascade involves activation by a dedicated E1 heterodimer followed by transfer to the SUMO E2 conjugating enzyme Ubc9. Ubc9 catalyzes SUMO transfer to the final acceptor lysine which often (but not always) resides within a consensus motif ΨKxE (Ψ=hydrophobic aminoacid), and is sufficient for SUMOylation *in vitro*^2–4^. Nevertheless, non-catalytic SUMO E3 ligases can facilitate Ubc9 or enable substrate specificity, and seem to be required for SUMOylation in some contexts and perhaps for non-consensus sites *in vivo*^5,6^. SUMO is primarily nuclear, and since its discovery has emerged as an important regulator of different nuclear processes (reviewed in ref.^7^) such as transcription factor activity, DNA repair, rRNA biogenesis, chromosome organization and segregation. Mechanistically, SUMOylation may lead to diverse consequences including changes in protein conformation or localization, masking or competing with other PTMs, and most famously, regulating protein-protein interactions. SUMO-mediated interactions typically involve an aliphatic stretch flanked by acidic amino acids (SUMO interacting motif, SIM) in the partner protein and although individually weak in their nature, multivalent SUMO-SIM interactions were proposed to stabilize large molecular complexes^8,9^ and promote the formation of phase-separated compartments such as PML (promyelocytic leukemia protein) bodies^10^. However, despite being implicated in a myriad of biological processes, our understanding of SUMO’s mechanistic role within different molecular and cellular contexts is far from complete.

Previous work in *Drosophila* implicated the SUMO pathway in the regulation of heterochromatin establishment and piRNA-mediated transposon silencing^11–14^. In germ cells, Piwi clade proteins (Piwi, Aub and Ago3 in flies) and Piwi-interacting small RNAs (piRNAs) cooperate in intimately linked processes that ensure transcriptional and post-transcriptional transposon silencing and continuous production of mature piRNAs (reviewed in ref.^15^). Mature piRNA production and post-transcriptional cleavage of transposon RNAs by Aub and Ago3 occur in a dedicated perinuclear structure, the nuage. Antisense piRNAs produced in the nuage also become loaded in Piwi, which then enters the nucleus to find transposon nascent RNA and enforce co-transcriptional silencing at target loci^16–18^. To date, SUMOylation is known to participate in the nuclear piRNA pathway in several ways: First, the SUMO E3 ligase Su(var)2-10 was found to interact with piRNA pathway and heterochromatin proteins and play an essential role in the recruitment of the enzymatic complex SetDB1/Wde that deposits the silencing epigenetic mark H3K9me3^12^, as well as with the MEP-1/Mi-2 chromatin remodeler complex^14^. Second, SUMOylation of Panoramix (Panx) – a co-repressor required for H3K9me3 deposition downstream of Piwi – was found to mediate its interaction with the general heterochromatin effector Sov^13^. In addition to silencing of transposons, Su(var)2-10 and SUMO were found to control H3K9me3 deposition at piRNA-independent genomic loci such as developmentally silenced tissue-specific genes^19^. The pervasive effect of SUMO depletion on global H3K9me3 levels at diverse classes of genomic targets suggests that SUMOylation is a critical process in the regulation of repressive chromatin integrated within different regulatory pathways. However, further understanding of the role of SUMOylation on chromatin requires knowledge of the full spectrum of SUMO targets.

We developed a proteomics approach that enables the identification of SUMOylated proteins with aminoacid-level site predictions from different tissues in the classic model for piRNA and heterochromatin studies *D. melanogaster*. Here, we report a comprehensive dataset of SUMO targets in the fly ovary, which represents the most extensive analysis of the SUMOylated complement in a metazoan reproductive tissue to date. Notably, we identified strong enrichment of heterochromatin factors among SUMOylated proteins, supporting the notion that SUMO plays a complex role in heterochromatin regulation that extends to multiple targets and protein complexes. Moreover, we find a striking enrichment of proteins specific to the piRNA pathway among SUMO targets, including the central effector of epigenetic silencing by piRNAs, Piwi, and several proteins that localize to the nucleus and nuage. We further validated the SUMOylation of selected piRNA pathway factors: Piwi, Panx, Spindle-E (Spn-E) and Maelstrom (Mael) in germ cells and showed that the modification of Mael and Spn-E, but not Panx, is Piwi-dependent, indicative of multiple SUMO roles in distinct steps of transposon silencing. Altogether, our findings point to a previously unappreciated multilayered role of SUMOylation in the piRNA pathway and heterochromatin regulation that provide important clues toward our understanding of the molecular mechanisms of genome regulation and transposon control.

## Results

### Establishing a system for the detection of SUMOylated proteins in Drosophila

Proteome-wide studies of PTMs have benefited from the development of methods and reagents that enable specific enrichment of modified proteins or peptides from total protein lysates. Basic methods for the enrichment of SUMO-modified proteins involve pulldown with anti-SUMO antibodies. However, this approach prohibits the use of stringent washing conditions and is therefore prone to high background. Furthermore, endogenous SUMO does not have trypsin cleavage sites close to its C-terminus, and trypsin digestion of SUMOylated proteins generates large, branched peptides from modified regions that are incompatible with conventional bottom-up proteomics (Fig. 1A). Accordingly, specific modified residues remain unknown and SUMOylation events can only be inferred indirectly from the abundance of other peptides from purified SUMOylated proteins.

**Figure 1.**
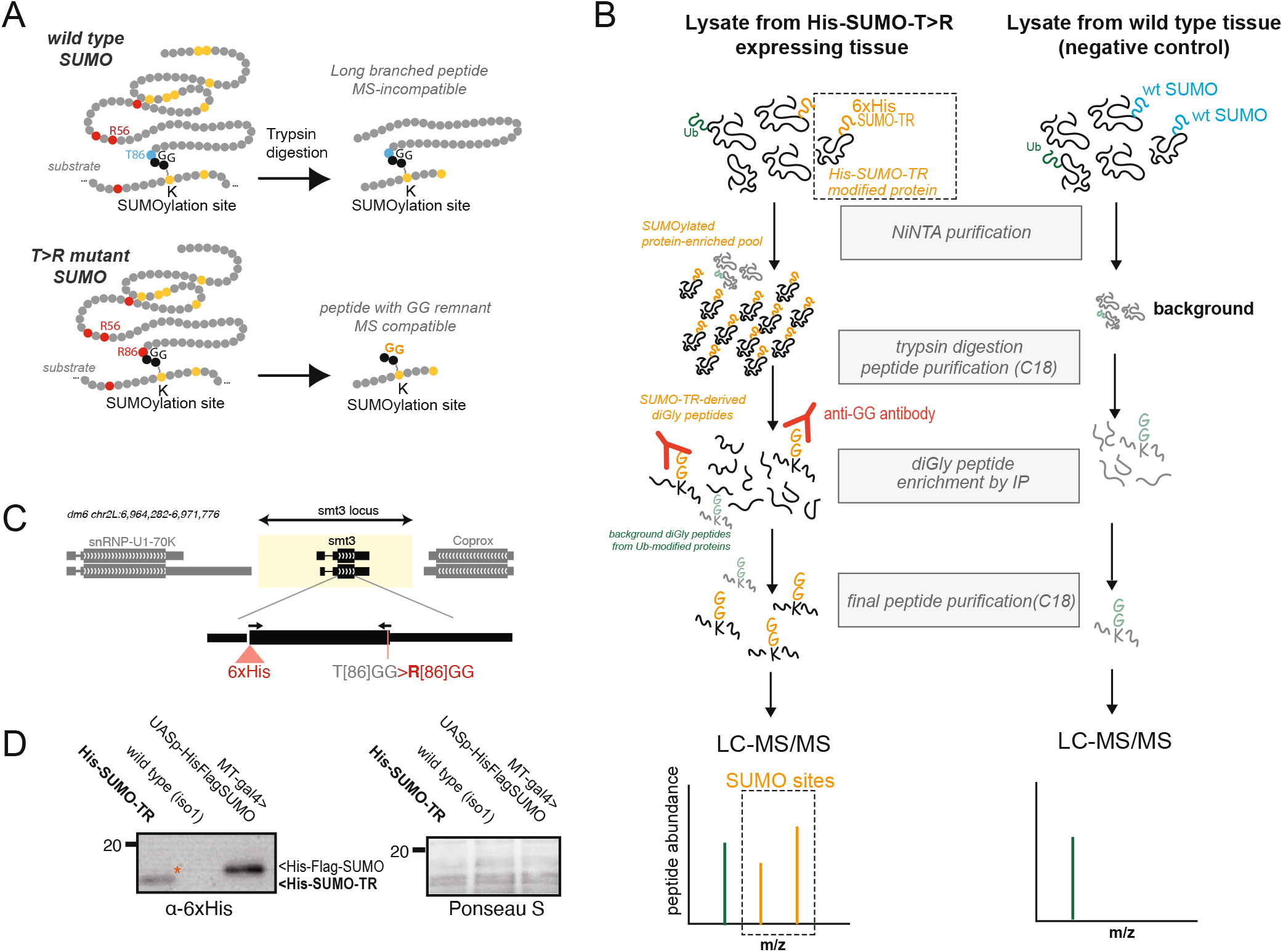
Proteomics approach for SUMO site detection. A. Diagram of the peptides generated after trypsin digestion of proteins modified by wild-type and T>R mutant SUMO. B. Diagram of the experimental design and workflow for SUMOylation-derived diGly-modified peptide MS analysis. C. Diagram of the recombinant SUMO construct used to generate SUMO-TR expressing transgenic flies. D. Western blot of total ovary lysate confirming the expression of recombinant SUMO. Ovaries from flies overexpressing 6xHisFlag-tagged SUMO are used as a positive control.

To overcome these obstacles and obtain a high-confidence dataset of the SUMO-modified proteins (herein referred to as the SUMOylome) in *Drosophila*, we adapted an approach that allows stringent purification, enrichment, and proteomic detection of peptides containing a modified SUMO remnant^20^. This approach, originally developed for the study of SUMOylation in human cells^20^, employs ectopically expressed SUMO protein with 6xHis-tag and a point mutation (T>R substitution, herein referred to as SUMO-TR) that enables trypsin cleavage before the C-terminal GG motif at the conjugation site (Fig. 1A,B). In a two-step purification process, SUMO-modified proteins are first selected based on the His tag by nickel affinity under denaturing conditions, which eliminates the activity of SUMO proteases and most background from noncovalently bound proteins. The resulting SUMOylated protein-enriched fraction is then trypsinized to generate a mixture of peptides including branched peptides with short di-glycine (diGly) remnant from cleaved SUMO-TR moiety. These diGly remnant-containing peptides are further selected from the total pool by pulldown with a specific antibody, purified, and analyzed by mass spectrometry. Of note, Ubiquitin and other ubiquitin-like protein modifiers naturally have an arginine residue before the terminal diGly motif, therefore, any ubiquitinated protein that unspecifically co-purifies with SUMOylated proteins during the His-based enrichment step can generate background diGly peptides. However, this background can be accounted for through negative control samples from tissues that do not express SUMO-TR (Fig. 1B). To enable this method for SUMOylation detection in *Drosophila* tissues, we created a transgenic line that carries a modified copy of the *smt3* locus (which encodes the single SUMO homolog in this species) where the SUMO coding sequence has a 6xHis N-terminal tag, T86>R substitution before the C-terminal GG motif, and a ~2.5kb upstream region containing the putative endogenous promoter. The His-tagged SUMO-TR protein was detectable by Western Blotting (Fig. 1C) and importantly, this transgene completely rescues the lethality of the null allele *smt3^04493^*, confirming that it encodes a fully functional protein.

### Identification and characterization of the Drosophila ovarian SUMOylome

We used flies expressing SUMO-TR to optimize the two-step purification procedure described above (Fig. 1B) and obtain SUMOylation site-derived peptides from ovarian tissue (See Materials and Methods for details). Following this procedure, we performed 3 independent experiments, where each replica involved parallel sample preparation from SUMO-TR ovaries and wild-type controls followed by label-free tandem mass spectrometry (MS). Each sample yielded ~1000-2500 diGly sites (Fig. 2A) and altogether, we detected 3159 exact diGly sites mapping to the protein products of 1295 genes. In comparison, MS analyses of input samples prior to diGly enrichment yielded less than 100 diGly sites (data not shown), highlighting the importance of the diGly enrichment step for capturing SUMOylation sites. For each diGly site, we calculated the normalized intensity ratios between SUMO-TR and control samples. The ratios showed a prominent bimodal distribution: approximately half sites in each experiment were detected exclusively in SUMO-TR samples, indicative of genuine SUMOylation targets (Fig. 2B). The diGly sites detected with similar intensities in SUMO-TR and control samples or biased to control samples likely originate from unspecifically bound ubiquitinated proteins. Consistent with this, motif analysis showed that diGly sites detected in SUMO-TR samples are enriched in the canonical SUMOylation motif, while diGly sites from the negative control do not have any motif enrichment (Fig. 2B, Fig. S1).

**Figure 2.**
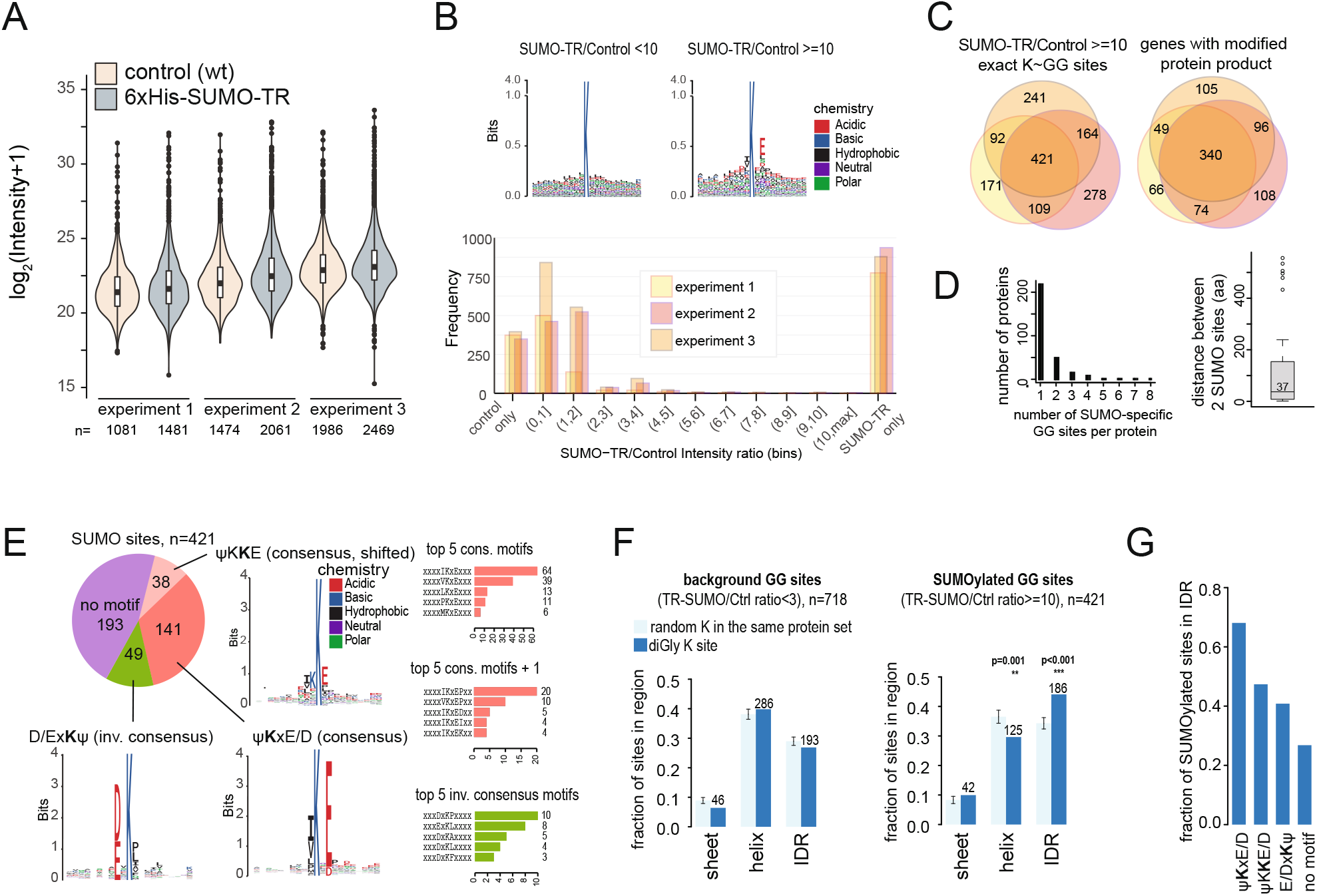
Characteristics of the ovarian SUMOylome. A. Distribution of diGly site intensities in different samples. B. (Top) Amino acid frequencies flanking predicted K-ε-GG, for sites with high and low SUMO-TR:control intensity ratios in a representative replicate. Also see Fig. S1. (Bottom) Distribution of SUMO-TR:Control intensity ratios of diGly sites in each experiment. C. Overlap of exact SUMO sites or SUMO-modified genes (irrespective of exact sites) between replicates. D. Numbers of predicted SUMO sites per protein and distance between sites for proteins with 2 sites. E. Sequence motifs at 421 high confidence SUMO sites. The predicted diGly-modified lysine is in bold. Bar graphs show the counts of topmost frequent aminoacids at positions −1 and +2, and −2,−1 and +2 from the modified lysine. F. Localization of diGly sites with high or low intensity ratios in predicted structured or intrinsically disordered regions, compared to randomly selected lysines from the same protein over 1000 iterations. G. Localization of diGly SUMO sites embedded in different motifs to IDRs.

The sets of predicted *bona fide* SUMO targets (SUMO-TR/Control intensity ratio >10) were highly reproducible between the 3 experiments, with ~50% overlap on the specific site level, and a 75% overlap on the target protein-coding gene level (Fig. 2C). Previously known SUMO targets including RanGAP1, PCNA, Su(var)2-10/dPIAS are present in the high confidence set. Specific sites that do not appear in all experiments tend to have the lowest intensities (Fig. S2). Considering that SUMOylation is often transient and affects a minuscule fraction of a given protein, it is likely that sensitivity is the major limitation to the reproducibility of detected sites between replicates. About ~25% of the proteins with diGly remnant have more than one modification site, with widely variable distances between two sites (Fig. 2D). This pattern could indicate that proteins can be alternatively SUMOylated at different residues, multi-SUMOylated, or even SUMO- and Ubiquitin-modified at the same time. As an aside, we detected several diGly sites on SUMO itself (Table S1); as some of these sites can be detected in the negative control, this pattern is indicative of hybrid SUMO-Ubiquitin chains.

To gain a further insight into the sequence features of the fly SUMOylome, we assessed the presence of amino acid motifs in the conservative set of 421 distinct high-confidence SUMOylation sites detected in all 3 experiments. Sequence pattern searches identified the canonical consensus and inverse consensus SUMOylation motifs (Ψ<u>K</u>xE/D and E/Dx<u>K</u>Ψ, respectively) at 45% of the sites (141 and 49) in this set (Fig. 2E). ~10% of the sites have the motif ΨK<u>K</u>E/D with the modification assigned to the second lysine – a pattern that might arise from uncertainty in diGly position prediction in the case of neighboring lysines. The remaining sites were not enriched for any known motif, and *de novo* motif search via MoMo did not identify any significant sequence bias around the central lysine (not shown), suggesting that as in other systems, the SUMOylation consensus motif is not obligatory. Like observations in human cells^17^, consensus sites displayed a marked preference for Isoleucine and Valine as hydrophobic residues. Additionally, a sizable fraction of the sites with strong SUMOylation consensus had a downstream Proline – an extended motif associated with SUMOylation/acetylation switch^21^ (Fig. 2E). Finally, we assessed the position of SUMOylation sites with respect to predicted structural features based on deep learning language models^22^ including alpha sheets, beta helices and intrinsically disordered regions (IDRs). This analysis demonstrated that SUMOylated sites are significantly overrepresented in IDRs and under-represented in structured regions compared to randomly selected lysines from the same protein or the background diGly sites (TR-SUMO/control ratio < 3 in all replicates). Similar results were obtained using IDR predictions generated with IUPred2A^23^ (Fig. S3). Notably, SUMOylation sites within canonical motifs showed the strongest bias towards IDRs (Fig. 2G).

Altogether, these results show that SUMOylation affects a large number of proteins in the *Drosophila* ovary, with the SUMOylome displaying conserved sequence and structural features with other organisms. Importantly, the breadth of the SUMOylome, together with the observations that a substantial fraction of SUMOylation sites do not have a specific association with a motif or structural features highlight the need for unbiased experimental approaches to identifying SUMOylation targets in different cellular and organismal contexts.

### Functional groups of SUMOylated proteins

To investigate the possible roles of SUMOylation in the ovary, we carried out functional enrichment and interactome analyses for the conservative set of proteins that carry reproducible SUMO sites in all experiments, i.e. 421 sites (Fig. 2C) in 292 proteins, using gene ontology (GO) and STRING (Fig. 3A,B, Table S2). Results showed that SUMOylation is common among specific physically and functionally linked groups of ribosomal and nuclear proteins (Fig. 3A,B, Table S2). Ribosomal proteins are highly abundant and hence a frequent source of background in proteomics. However, the number of reproducible diGly sites mapping to ribosomal proteins in the SUMO-TR samples is significantly above the background (ribosomal proteins carry 57 out of the 421 *bona fide* SUMO sites (13%), versus 37 out of the 714 background diGly sites(5%)) and could therefore represent genuine targets. Indeed, the SUMOylation machinery has been implicated in ribosomal protein regulation^24^. Beyond the ribosome, SUMO targets are common among proteins participating in rRNA processing, mRNA splicing, DNA damage response and repair, consistent with a conserved role of the SUMO pathway in these processes from yeast to mammals (reviewed in ref.^7^). Notably, SUMOylation is widespread among chromatin-associated proteins. High-confidence SUMOylation sites can be found in several core histones and the HP1 paralogs HP1b and HP1c; a weaker site was also detected in the central heterochromatin effector HP1a/Su(var)205 at position homologous to the SUMO site in its mammalian homolog HP1ɑ/CBX5 (K32). Strikingly, among the most enriched gene ontology terms in the SUMOylated set are terms related to heterochromatin and transposon repression; for example, 21 out of the 96 genes associated with the “heterochromatin organization” GO term, and 8 out of 25 genes associated with the “negative regulation of transposition” GO term are in the high confidence SUMO target set.

**Figure 3.**
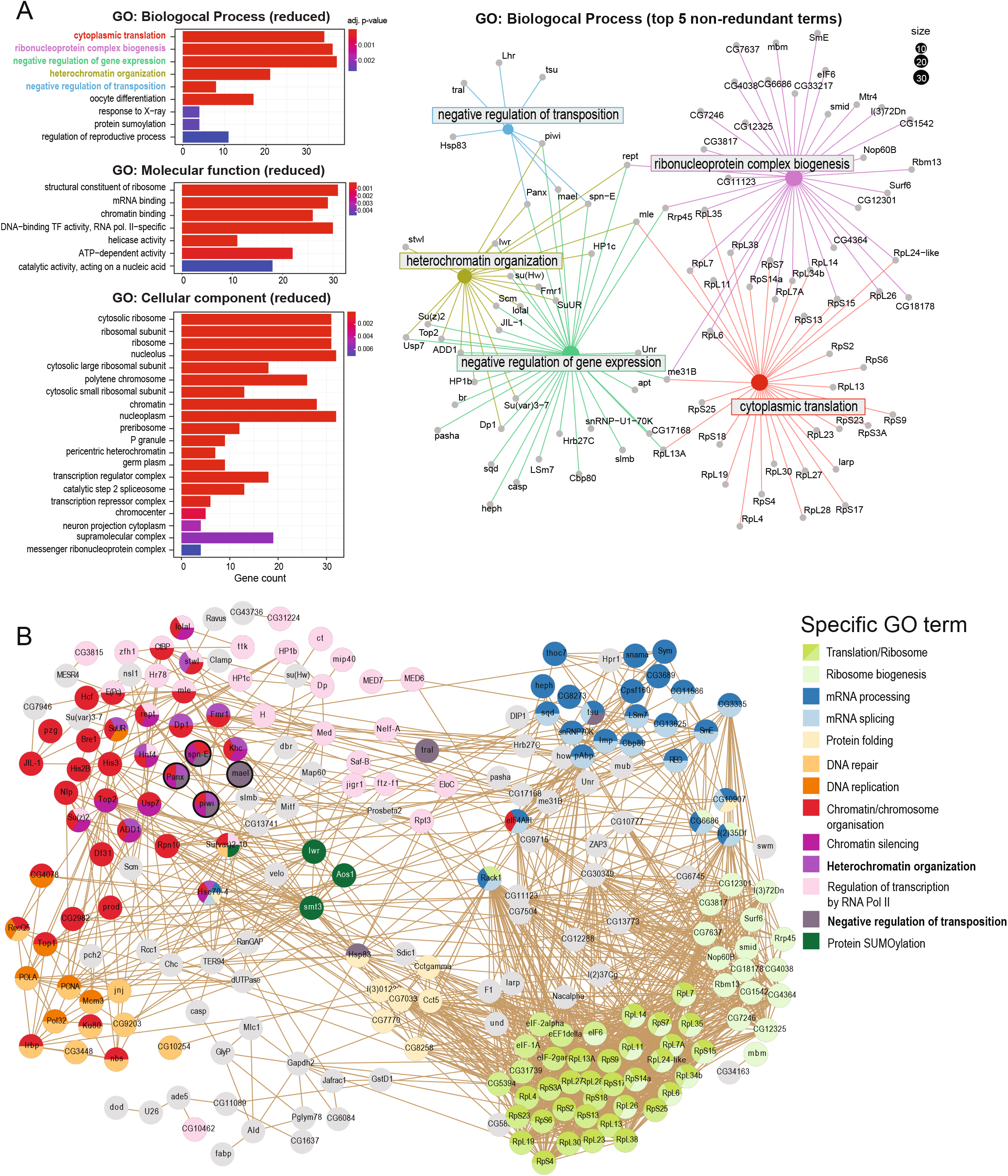
Functional groups of SUMOylated proteins. A. Summary of Gene Ontology analysis. (Left) Bar charts of the most enriched GO terms and corresponding gene numbers among the high confidence SUMOylated targets after semantic simplification, ordered by adjusted p-value. (Right) Network diagram of proteins annotated in the top 5 enriched Biological Process GO categories. B. STRING diagram of the interactions between identified high confidence SUMOylation targets. Single nodes are not shown. Nodes are colored according to selected GO terms and manually grouped according to functional annotations.

To further examine the breadth of SUMOylation within heterochromatin- and piRNA pathway-associated factors that control transposons in the ovary, we mapped the presence of SUMOylation sites among the physical interactors of the key heterochromatin effectors SetDB1/Egg, Wde and HP1a, as well as nuclear piRNA pathway effector Piwi, and Vasa – central component of the nuage compartment where piRNA biogenesis and transposon RNAi-mediated cleavage take place^25–27^ (Fig. 4A). Several HP1a interactors are modified by SUMO, including the split orthologs of the mammalian chromatin remodeler ATRX dADD1 and XNP, pointing to a multifaceted role of SUMOylation in transcriptional silencing and heterochromatin regulation. Furthermore, several high confidence SUMOylation sites are present in Piwi, as well as several others piRNA pathway proteins including Panoramix (Panx) – a component of the so-called SFiNX/Pandas/PICTS^28–30^ complex required for H3K9me3 deposition downstream of Piwi (also recently reported as SUMO target in another study^13^), Maelstrom (Mael) – essential for transcriptional repression and piRNA biogenesis from dual-stranded clusters^16,31^, and the nuage component Spindle-E (SpnE) – a germline-specific DEAD-box helicase essential for piRNA production^32–35^. Additionally, manual inspection of the data showed that several proteins involved in piRNA biogenesis such as thoc7, CG13741/Bootlegger and Hel25E can be SUMOylated (Table S1). The presence of SUMO at a wide range of proteins involved in piRNA biogenesis and function suggests that this modification may regulate the cellular response to transposon activity at multiple levels including but not limited to previously identified SUMO functions.

**Figure 4.**
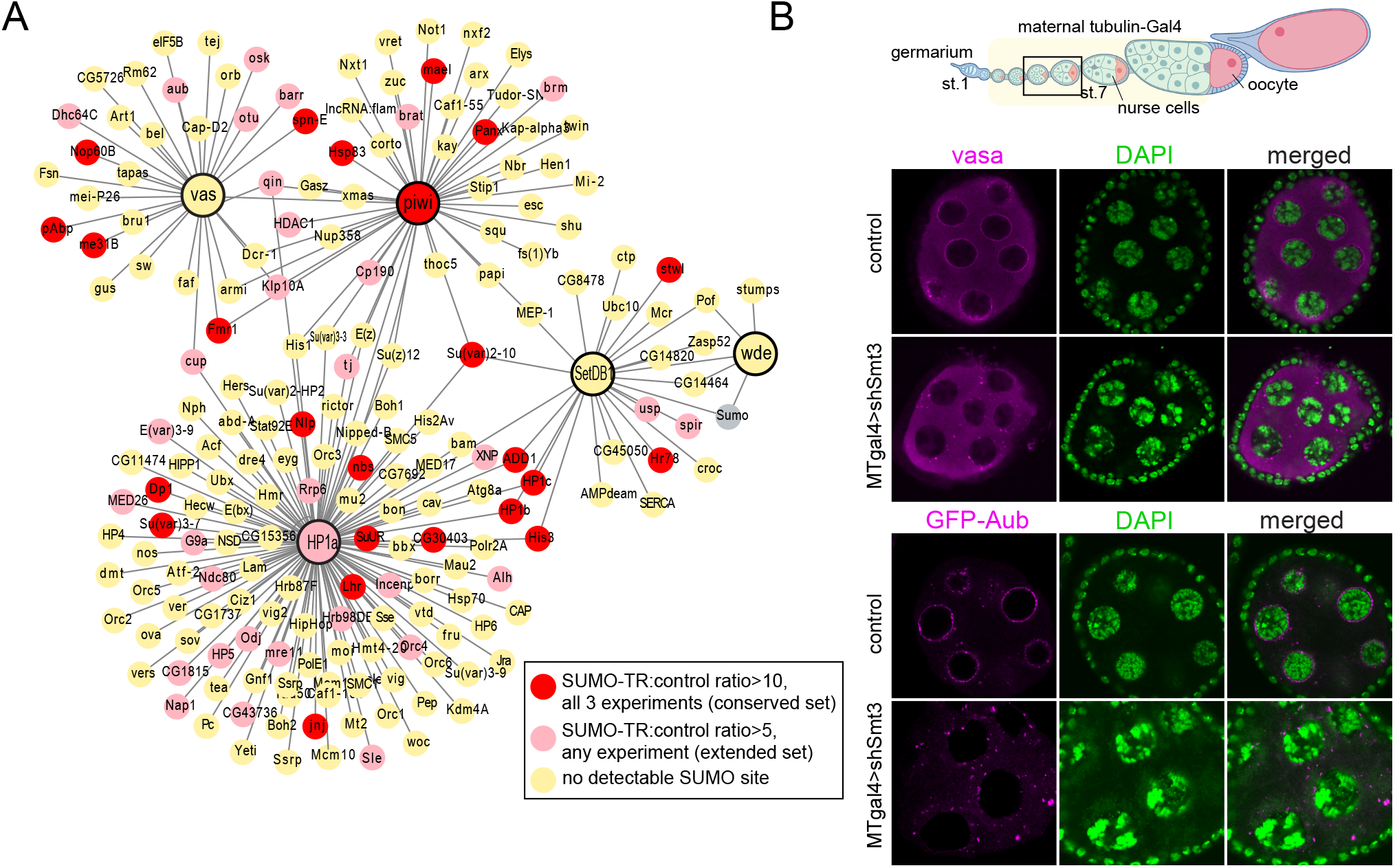
SUMO targets nuclear and cytoplasmic piRNA factors. A. Network diagram showing physical interacting partners of Piwi, HP1, SetDB1, Wde, and Vasa proteins retrieved from FlyBase. Nodes are colored according to detection frequency of SUMOylation sites in the MS data. Additional interactions between partner nodes are not shown. B. (Top) Diagram of an ovariole with indicated stages where the maternal tubulin-Gal4 (MT-gal4) driver is active; stages of imaging are highlighted. (Bottom) Confocal images of Vasa (immunostaining) and Aub (GFP-tagged Aub) localization in stage 5-7 egg chambers from flies expressing SUMO shRNA under the control of the MT-gal4 driver, or no driver control siblings.

Following the observation that several nuage-associated proteins are SUMO-modified, we wondered if the SUMO pathway affects the cytoplasmic piRNA compartment in addition to its previous implication in the nuclear piRNA pathway^13,19^. To explore this possibility, we investigated the subcellular localization of the core nuage component Vasa, and the nuage-localized piRNA effector Aub, upon SUMO depletion. To this end, we employed a previously characterized small hairpin RNA targeting SUMO^12^ under the control of maternal tubulin Gal4 driver, which is active from stage 2-3 of oogenesis onward. In this condition, oogenesis fails to complete with nurse cell nuclei collapsing around stage 7, however, prior to that egg chambers appear morphologically normal and can be analyzed (Fig. S4). As previously reported, under normal conditions Aub and Vasa appear as a relatively smooth perinuclear layer^36^ (Fig. 4). However, SUMO knockdown leads to markedly disrupted Aub and Vasa localization at the nuclear periphery and dispersed granules throughout the cytoplasm (Fig. 4B). Collectively, these results indicate that the SUMO pathway may support transposon silencing not only through its involvement in heterochromatin formation but also through effects on the nuage compartment.

### SUMOylation of piRNA pathway factors in the female germline

To further understand SUMO’s roles in the piRNA pathway, we sought to validate the SUMOylation of four essential piRNA pathway proteins – Piwi, Panx, Spn-E and Mael. As SUMOylation typically affects only a small fraction of the total protein pool, and the detection of such species is limited by antibody availability and affinities, we devised a sensitive system to analyze SUMOylation of proteins of interest in the germ cells of the ovary which express the full cytoplasmic and nuclear piRNA pathway. Specifically, we utilized UASp/Gal4 to express Flag-tagged SUMO and GFP-tagged target protein in nurse cells from stage 2-3 of oogenesis and later using the maternal tubulin-Gal4 driver (except GFP-Piwi which was under the control of its native regulatory region). In this system, the GFP tag and high affinity anti-GFP nanobody allows protein purification under stringent washing conditions, while the sensitive and specific monoclonal anti-Flag antibody maximizes detection of small amounts of SUMO modified proteins by Western Blotting (WB).

Analyses of immunopurified GFP-Spn-E, Piwi, Panx and Mael from ovaries showed Flag-SUMO conjugated higher molecular weight forms consistent with SUMO modification in all cases (Fig. 5A). Note that free Flag-tagged SUMO migrates at about ~17 kDa on WB, slightly higher than its predicted molecular weight of ~11 kDa. Each target displayed multiple higher molecular weight bands, supporting the presence of multiple SUMO moieties or mixtures of SUMO and other protein PTMs on the same protein. The migration pattern of Panx indicates the existence of mono-SUMOylated and an array of poly-modified forms. A similar pattern of multiple modified Panx species was reported in another recent study where authors used a custom-made antibody against the native protein^13^. The observed shift in molecular weight for GFP-Piwi from ~120kDa to ~160kDa and above indicates that it carries two or more protein modifiers, at least one of which is SUMO. Mael’s and SpnE’s SUMO-modified forms have molecular weights of ~40-50 kDa greater than their unmodified forms, also consistent with at least 2-3 modified sites within the same protein. Of note, the number of higher molecular weight bands in Mael and Panx exceeds the number of predicted diGly sites. While our proteomics detection is limited to peptides within a particular size range and thus is unlikely to cover all possible modified residues, we also cannot rule out the existence of poly-SUMO or hybrid SUMO-Ubiquitin chains (see Discussion).

**Figure 5.**
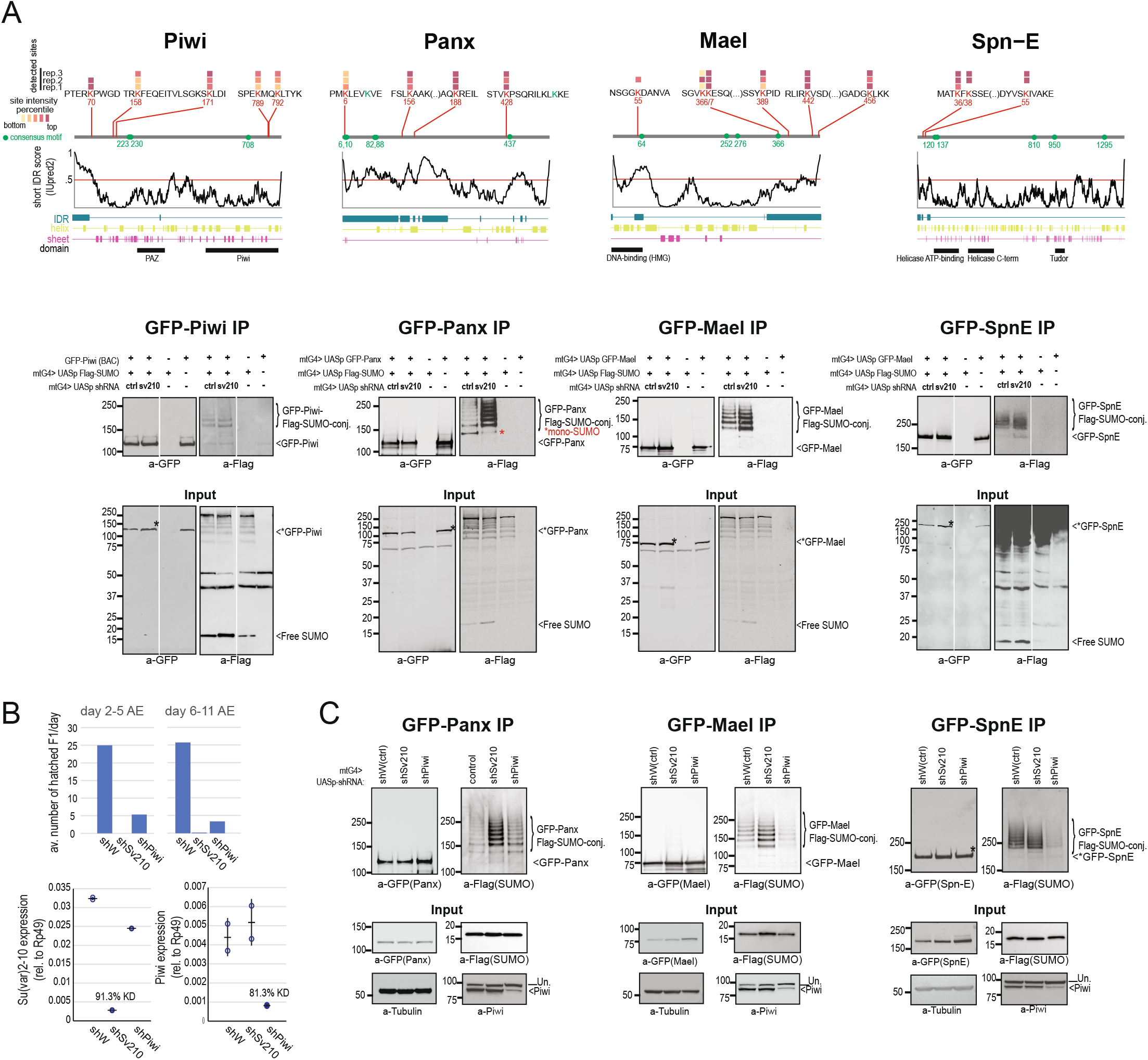
SUMOylation of Piwi, Mael, Spn-E and Panx. A. (Top) Diagrams of structural features, consensus SUMOylation motifs, and experimentally detected diGly sites with TR-SUMO:Control intensity ratios >10. Color indicates diGly site intensity percentiles. (Bottom) WB analysis of target SUMOylation in the female germline. Ovary lysates from flies carrying indicated transgenes (mtG4=maternal tubulin Gal4 driver) were subjected to GFP IP followed by WB detection first with anti-Flag antibody, and after stripping, anti-GFP antibody. Note that SUMOylated species are only detectable by the highly sensitive anti-Flag antibody, but not anti-GFP antibody. Lanes in Piwi and Spn-E gels were cropped for sample order consistency. Images are representative of 3-4 independent experiments per target. B. Efficiency of *su(var)2-10* and *piwi* germline knockdown estimated by fertility test (top) and RT-qPCR (bottom). Results are from samples from the same genetic crosses used for Western blotting in C, GFP-Panx. C. WB of target SUMOylation in the female germline upon *white* (control), *su(var)2-10* and *piwi* germline knockdown. Images are representative of two independent experiments. Un.=unspecific band.

We also tested whether ectopically overexpressed piRNA pathway proteins can become SUMOylated in S2 cells, a somatic cell line from embryonic haematocyte origin which does not have an active piRNA pathway. While SUMOylation was still detectable, its pattern was drastically different from the characteristic “ladder” observed in the female germline (Fig. S5). The tissue-specific patterns of Piwi, Panx, Mael and SpnE SUMOylation therefore suggest that this modification is involved in specific regulatory steps in the germline in the context of active piRNA pathway response to genomic parasites.

Previous work identified the SUMO E3 ligase Su(var)2-10 as an effector of transposon silencing and heterochromatin establishment that acts downstream of Piwi^12^. To test if Su(var)2-10 regulates the SUMOylation of Mael, Panx, Spn-E or Piwi, we induced its germline-specific knockdown via previously characterized UASp-controlled small hairpin RNA (shRNA)^12^. Su(var)2-10 depletion did not abolish the SUMOylation of any of the investigated proteins, despite efficient knockdown evidenced by female sterility and RT-qPCR (Fig. 5B). In fact, we systematically observed (n=2-4 independent experiments for each protein) a slight increase of SUMOylated species upon Su(var)2-10 germline knockdown. Of these, the most prominent change was seen for Panx, which displayed a marked increase of its poly-modified forms and decrease of mono-SUMOylated form (Fig. 5A, asterisk). Thus, it appears that Su(var)2-10 is not a SUMO E3 ligase for Piwi, Mael, Spn-E, or Panx, but the modification of these proteins in germ cells is regulated in a manner influenced by Su(var)2-10 loss.

As Su(var)2-10 loss leads to strong transposon upregulation^12^, the increased SUMOylation of piRNA pathway proteins upon Su(var)2-10 knockdown may indicate SUMO implication in protein complexes actively engaged in transposon response. To explore this possibility, we decided to analyze the SUMOylation of Panx, Spn-E, and Mael upon Piwi depletion that disrupts the nuclear piRNA-dependent silencing machinery. To this end, we utilized UASp-controlled shRNA against Piwi, which induces efficient Piwi germline knockdown (Fig. 5B,C). Notably, Piwi depletion resulted in marked loss of Mael and Spn-E SUMOylated forms (Fig. 5C), suggesting that the modification of both proteins in germ cells depends on Piwi. In contrast, Piwi germline knockdown did not affect the number and levels of SUMO-modified Panx forms, indicating that Panx SUMOylation is governed by a distinct mechanism independent of the piRNA-Piwi complex in this cellular context. Altogether, these results indicate that SUMOylation of Mael and Spn-E occur in a regulated manner downstream of Piwi, pointing to novel and multifaceted roles of SUMOylation in the piRNA pathway.

## Discussion

The SUMO pathway is essential for normal cell function as evidenced by the severe phenotypes of loss-of-function mutants of SUMO and SUMO ligases in various systems^37^. Our mechanistic understanding of SUMOylation however has remained limited, partly owing to the technical difficulties of detecting this modification. Here, we adapted the diGly remnant enrichment method for unbiased discovery of SUMO-modified proteins with specific site predictions in *Drosophila*. We note that while diGly peptide enrichment improves the sensitivity and specificity of SUMOylated protein detection, as with other bottom-up proteomics methods, detection is only possible for sites residing in protease-generated peptides within a specific size range. Also, exact site assignment might be imprecise when two lysine residues are in proximity, and in the event a protein simultaneously carries SUMO and another modification that leaves a diGly remnant after trypsin cleavage such as ubiquitin (in such cases, the target protein would be enriched in the SUMOylated pool but detected diGly remnants might come from any concomitant modification). Despite these limitations, the absolute identification of SUMO protein targets remains of high confidence, and predicted sites are informative to narrow down exact modified residues for future mutational studies. Altogether, our proteomics survey of the *Drosophila* ovary uncovered a large SUMOylated complement that displays conserved features of this modification, including sequence and structural biases of preferred SUMOylation sites, collective targeting of proteins from the same process/complex, and enrichment of SUMO targets among proteins involved in various aspects of nucleic acid metabolism. Notably, we found SUMO targets with complex modification patterns among a variety of piRNA pathway proteins functioning in both transcriptional and post-transcriptional transposon silencing, pointing to a multifaceted role of this modification in the cellular response to genomic parasites.

### Complexity of the SUMO-modified proteome

Approximately a third of all SUMO targets in our high confidence set contained two or more diGly remnants, and analysis of selected proteins in germ cells by Western blotting (Fig. 5) demonstrated a characteristic “ladder” of multiple modified forms. These patterns are indicative of multi-SUMOylation (when multiple residues in the same protein are SUMOylated), and possibly, concomitant modification by SUMO and other protein modifiers. In addition to multiple modifications at different residues, studies in yeast and mammalian systems have identified a variety of homotypic and heterotypic SUMO and Ubiquitin chains, as well as hybrid SUMO-Ubiquitin chains that could create a complex regulatory “code”^38,39^. We detected several diGly sites on the endogenous SUMO proteins in control samples (Table S1), confirming the presence of hybrid SUMO-Ubiquitin chains in various configurations in flies. *In vitro* studies and comparative genomics suggested that SUMO-only chains – common in yeast, plants and mammals – have been evolutionarily lost in flies^40^. Distinguishing SUMO chains from SUMO-Ubiquitin hybrid chains is not possible with our approach. However, considering the presence of diGly remnants at syntenic positions to typical sites of SUMO chain formation in other species, the existence of SUMO chains *in vivo* merits future investigation.

### Heterochromatin as a SUMOylation ‘hot-spot’

A prominent feature of the identified SUMO targets is that they are often found among physically and functionally related proteins. This is consistent with previous notions of an unusual property of SUMOylation compared to other PTMs, namely, that SUMO ligases can modify entire groups of physically interacting proteins at multiple and perhaps redundant sites (reviewed in ref.^9^). Such group SUMOylation is thought to create multiple SUMO-SIM interactions within large molecular complexes that act synergistically to facilitate their assembly and function. Previous examples of “SUMO hot spots” include DNA repair pathway effectors in yeast, ribosome biogenesis, or PML bodies^9^. The SUMO pathway has long been linked to heterochromatin: SUMOylated histones, SUMO and SUMO ligases were shown to localize to heterochromatic regions in various systems, and several essential heterochromatin proteins including HP1, H3K9 methyltransferase and histone deacetylase complex components were identified as SUMO targets or interactors (reviewed in ref.^41^). Our data identifying dozens of heterochromatin proteins as SUMOylation targets suggests that heterochromatin can be considered another hotspot of group SUMOylation. Moreover, SUMO was recently shown to affect interconnected repressive chromatin factors in mouse embryonic stem cells^42^, pointing to an evolutionary conserved role of SUMOylation in the regulation of chromatin organization. Notably, SUMO polymers were shown to drive phase separation in different subcellular contexts^10,43^. In the future, it will be interesting to establish the functional significance of the heterochromatic SUMO target spectra, particularly from the perspective of the biophysical properties of heterochromatin and heterochromatin-related multi-subunit regulatory complexes.

### A multifaceted role of SUMO in the piRNA pathway

The discovery of a large set of SUMO-modified proteins among piRNA pathway factors paints a complex picture of the roles of SUMOylation in the cellular response to transposons. First hints on the implication of SUMO in the piRNA pathway emerged from genetic screens to identify factors required for transposon repression^11^. More recently, functional studies demonstrated that SUMO is essential for the nuclear compartment of the piRNA pathway that enforces co-transcriptional silencing of transposons by the installation of the silencing mark H3K9me3 at target loci. First, transcriptional silencing was shown to involve Su(var)2-10/SUMO-dependent recruitment of chromatin modifying complexes downstream of Piwi and the co-repressor Panx^12,14^. Additionally, transposon silencing requires a SUMO-mediated interaction between Panx and the general heterochromatin factor Sov^13^. The proteomic identification of numerous general heterochromatin proteins as well as Panx itself as SUMO targets emphasizes the possibility that SUMOylation plays a multifaceted role in the transcriptional silencing of transposons by piRNAs. However, in addition to that, we also uncovered SUMO targets among piRNA proteins which belong to spatially and functionally distinct piRNA pathway compartments such as the nuage, and found that SUMO depletion disrupts the nuage structure, suggesting that SUMOylation has even broader implications in transposon silencing.

While dissecting the multiple roles of SUMOylation in the piRNA pathway will require extensive future work, observations of the SUMOylation patterns of selected piRNA pathway proteins provide several interesting clues. First, the SUMO-modified forms of all four proteins are of relatively low abundance and only detectable by highly sensitive antibodies, indicative of transient nature. Dynamic and reversible combinatorial modifications via SUMO and SUMO-Ubiquitin chains can create unique surfaces for multivalent protein interactions. Complex chains were previously shown to act as signals in the regulated recruitment and assembly of protein complexes involved in centromere organization and DNA damage repair (reviewed in ref.^38^). It is possible that analogously, complex SUMOylation patterns may be involved in dynamically regulated steps of piRNA biogenesis and function. A second line of clues comes from the different effects of Piwi and Su(var)2-10 germline knockdown on the SUMOylation of the examined piRNA pathway proteins. The SUMO E3 ligase Su(var)2-10 is essential for H3K9me3 deposition at transposon targets^12^. The finding that Piwi, Panx, Spn-E or Mael SUMOylation do not depend on Su(var)2-10 is consistent with a model where Su(var)2-10 acts at downstream steps of heterochromatin establishment, but SUMOylation is also involved in upstream or parallel processes related to piRNA biogenesis and function. Moreover, as the SUMO-modified forms of the four proteins increase upon Su(var)2-10 knockdown, it is tempting to speculate that SUMOylation might be associated with increased piRNA pathway activity in response to the strong transposon upregulation that occurs upon this genetic perturbation. Nevertheless, it is important to consider that Su(var)2-10 depletion causes substantial transcriptomic changes beyond transposon activation, including the up- and down-regulation of hundreds of protein coding genes^12,19^, with possible indirect consequences on the SUMOylome.

Interestingly, Piwi depletion has differential effects on Mael, Spn-E, and Panx SUMOylation: Mael and Spn-E lose their SUMOylation in Piwi knockdown germ cells, while Panx SUMOylation remains unaffected (Fig. 5C). Previous studies recognized Mael as an essential factor for transcriptional repression at piRNA targets as well as piRNA-independent genomic loci, although its precise mechanistic role has remained unclear^16,31^. Considering the nuclear functions of Mael, loss of its SUMOylation upon Piwi knockdown might reflect a regulatory step following piRNA/Piwi recognizing their targets. Nevertheless, Mael also localizes to the nuage^16^, and further work will be required to establish whether Mael SUMOylation is related to its roles in the nucleus or a yet unknown function in the nuage. A possible role of SUMOylation in the cytoplasmic piRNA pathway branch is further highlighted by the modification of Spn-E. Spn-E is a putative RNA helicase which, despite unclear mechanistic role, is a well-established nuage component essential for piRNA biogenesis^32–35^. Since Piwi operates in the nucleus, yet its loading with piRNA occurs in the nuage, the dependence of Spn-E SUMOylation on Piwi raises the intriguing possibility that SUMO-mediated interactions may be involved in certain aspects of piRNA biogenesis. Future work combining our highly sensitive and robust germline SUMOylation assay with different genetic perturbations and SUMOylation-deficient mutants promises to improve our understanding of the precise mechanisms that orchestrate the piRNA pathway in its different subcellular contexts.

Finally, the lack of effect of Piwi loss on the SUMOylation of Panx – one of the core components of the RNA-binding SFiNX/Pandas/PITSC complex that is essential for H3K9me3 deposition downstream of Piwi – suggests Panx is modified in a process that occurs independently of Piwi-mediated target recognition. Surprisingly, this result contrasts with recent data from OSC cells – an immortalized cell line derived from the ovarian somatic follicle cells which expresses only the nuclear portion of the piRNA pathway – where Panx SUMOylation was found to completely depend on Piwi^13^. While we cannot rule out that residual levels of Piwi may be sufficient to maintain Panx SUMOylation in our system, it is also possible that the regulation of Panx differs between the germline and soma, or even at different stages of oogenesis. A potential mechanistic difference in the nuclear piRNA pathway between germline and soma has critical implications to our understanding of this process and will be important to address in the future.

## Materials and Methods

### Drosophila stocks and husbandry

Drosophila lines encoding GFP-tagged Piwi under the native promoter (3rd chromosome), UASp-GFP -Panoramix, -Maelstrom, and -Spn-E, on the 2nd chromosome, and small hairpin RNA against the *smt3* gene, and GFP-tagged Aub under the control of the endogenous Aub promoter, were previously generated in Aravin/Fejes Toth laboratories^12,18,36^. Flies encoding UASp-driven HisFlag-tagged SUMO on the 3rd chromosome were a gift from Courey Laboratory^44^. Flies encoding shRNA against *su(var)2-10*, *piwi*, and *white*, the maternal tubulin(Mt)-Gal4 driver, the iso-1 Celera sequencing strain, and smt[04493] were obtained from the Drosophila Bloomington stock center (#32956, #33724, #33623, #7063, #2057, #11378). To study SUMOylation in germ cells, four transgenes including UASp-HisFlag-SUMO, GFP-tagged target, UASp-shRNA and Mt-Gal4 driver were combined by crossing balanced parental lines encoding GFP-tagged target and Mt-Gal4 driver, and lines carrying UASp-shRNA and UASp-HisFlag-SUMO.

To generate SUMO-TR flies, the smt3 locus + ~2kb upstream region fragments (chr2R:21704005-21717574, dm6 reference genome) were amplified with the introduction of a 6xHis tag and a T86>R mutation using Gibson assembly, subcloned into a phiC31 backbone based on the pCasper5 vector, and integrated into the P{CaryP}attP2 locus at BestGene Inc.

All stocks were maintained at 25°C on standard molasses-based media. Experiments were performed on samples from 3–5-day old females maintained on standard media supplemented with yeast for 2 days prior to dissections. Ovaries were manually dissected in cold PBS, flash frozen in liquid nitrogen and stored at −80°C.

### Proteomics sample preparation

The preparation of SUMO-derived GG-modified peptide samples was adapted from Impens et al. with some modifications^20^. All procedures were performed using low binding plasticware, HPLC grade water, and freshly prepared solutions. Hand dissected ovaries from 600 2-5-day old *D.melanogaster* individuals of smt3[04493]/CyO; {6xHis-smt3[T86R]}attP2 genotype (SUMO-TR) or iso-1 (Bloomington #2057) (control) were used in each experiment. Frozen samples were immediately lysed in 2.5 mL denaturing buffer (6M guanidium HCl, 10 mM Tris, 100 mM sodium phosphate buffer pH 8.0) using glass Potter-Elvehjem tissue grinder. Lysates were cleared by centrifugation at 20,000rpm for 10 minutes at 4°C. Cleared lysates were treated by 5mM tris(2-carboxyethyl)phosphine for 30 min at 37°C with gentle rotation, followed by 10 mM N-ethylmaleimide (Sigma) for 30 min at room temperature, and finally, 10 mM Dithiothreitol (Sigma). Lysate volume was brought to 8 mL with lysis buffer and imidazole was added to 5 mM final concentration. Lysates were incubated with 1 mL HisPur™ Ni-NTA Resin (ThermoFisher) overnight at 4°C with gentle rotation. After incubation, the Ni-NTA slurry was washed once with lysis buffer supplemented with 10 mM imidazole, once with wash buffer 1 (8M urea, 10 mM Tris, 100 mM sodium phosphate (pH 8.0), 0.1% Triton X-100, 10 mM imidazole), and three times with wash buffer 2 (8 M urea, 10mM Tris, 1 mM sodium phosphate buffer pH 6.3, 0.1% Triton X-100, 10mM imidazole), followed by two rounds of elution with elution buffer (300mM imidazole, sodium phosphate buffer pH 6.8) for 2 hours at 4°C to a final eluate volume of 1.5 mL. The eluate volume was increased to 10 mL with 50 mM ammonium bicarbonate and treated with sequencing grade trypsin (Promega, V5111) at 1:50 trypsin:protein ratio overnight at 37°C with gentle agitation. The following steps were performed according to the manual of the PTMScan® HS Ubiquitin/SUMO Remnant Motif (K-ε-GG) Kit (Cell Signaling #59322). In brief, trypsinized samples were acidified to pH 2-3 by the addition of 0.5 mL 20% trifluoroacetic acid (TFA), kept on ice for 15 mins, and centrifuged to remove potential precipitates. Peptides were then purified on a C18 column according to the kit manual, and lyophilized for >24hrs. Dry peptides were then resuspended in an IAP buffer (Cell Signalling #59322) and incubated with anti-diGly antibody-conjugated slurry (Cell Signaling #59322) at 4°C overnight, followed by two washes with IAP buffer, and 3 washes with HPLC-grade water. Finally, peptides were eluted with 0.15% TFA, purified using C18 StageTips, vacuum dried, and submitted to the Caltech Proteome Exploration Laboratory for Mass spectrometry analysis.

### LC-MS/MS and raw data processing

Label-free peptide samples were subjected to LC-MS/MS analysis on an EASY-nLC 1200 (Thermo Fisher Scientific, San Jose, CA) coupled to a Q Exactive HF Orbitrap mass spectrometer (Thermo Fisher Scientific, Bremen, Germany) equipped with a Nanospray Flex ion source. Samples were directly loaded onto an Aurora 25cm x 75μm ID, 1.6μm C18 column (Ion Opticks) heated to 50°C. The peptides were separated with a 60 min gradient at a flow rate of 220 nL/min for the in-house packed column or 350 nL/min for the Aurora column. The gradient was as follows: 2–6% Solvent B (3.5 min), 6-25% B (42 min), 25-40% B (14.5 min), 40-98% B (1 min), and held at 100% B (12 min). Solvent A consisted of 97.8 % H2O, 2% ACN, and 0.2% formic acid and solvent B consisted of 19.8% H2O, 80 % ACN, and 0.2% formic acid. The Q Exactive HF Orbitrap was operated in data dependent mode with the Tune (version 2.7 SP1build 2659) instrument control software. Spray voltage was set to 1.5 kV, S-lens RF level at 50, and heated capillary at 275°C. Full scan resolution was set to 60,000 at m/z 200. Full scan target was 3 × 106 with a maximum injection time of 15 ms. Mass range was set to 400−1650 m/z. For data dependent MS2 scans the loop count was 12, AGC target was set at 1 × 105, and intensity threshold was kept at 1 × 105. Isolation width was set at 1.2 m/z and a fixed first mass of 100 was used. Normalized collision energy was set at 28. Peptide match was set to off, and isotope exclusion was on. Data acquisition was controlled by Xcalibur (4.0.27.13), with ms1 data acquisition in profile mode and ms2 data acquisition in centroid mode.

Thermo raw files were processed and searched using MaxQuant (v. 1.6.10.43)^45,46^. Spectra were searched against *D. melanogaster* UniProt entries plus the His-SUMO-TR sequence and a common contaminant database. Trypsin was specified as the digestion enzyme and up to two missed cleavages were allowed. False discovery rates were estimated using a target-decoy approach, where the decoy database was generated by reversing the target database sequences. Protein, peptide, and PSM scores were set to achieve a 1% FDR at each level. Carbamidomethylation of cysteine was specified as a fixed modification. Protein N-terminal acetylation, methionine oxidation, and diGly remnant on lysine were specified as variable modifications with a maximum of 5 mods per peptide. Precursor mass tolerance was 4.5 ppm after mass recalibration and fragment ion tolerance was 20 ppm. Search type was specified as Standard with multiplicity of 1. Fast LFQ and normalization were used, and both re-quantify and match-between-runs were enabled.

### Bioinformatics analysis of SUMOylation sites

Summary tables of the normalized diGly sites output from MaxQuant (GlyGly (K)Sites.txt, Table S1) were analyzed and figures were generated using custom R scripts (*associated R markdown notebooks will be available from GitHub after peer review*). In brief, SUMO sites were considered sites with MS intensity ratios in the SUMO-TR to corresponding control samples >10, and background diGly sites were considered sites with SUMO-TR/control ratios <3. Motif searches were performed using regular expressions and the MoMo suite^47^ using a 11 aminoacid window centered on the predicted GG-modified lysine. Protein structure predictions were performed using the ‘bio_embeddings’ package with default parameters^22^, or IUPred2 with default parameters^23^. To test whether the numbers of diGly sites falling within a specific structural region are different than expected by random chance, we created a bootstrap distribution (n=1000) of the fraction of randomly selected lysines from the same set of proteins falling within each region. Functional enrichment analyses presented on Figure 3 were carried out using the clusterProfiler R package with annotations from the OrgDb package (GO source date 2021-Sep13). Semantic simplification was used to merge highly redundant terms. We note that similar results were obtained using TopGO, STRING and the bINGO Cytoscape plugin (not shown). STRING Interactions were plotted using CytoScape at 0.6 confidence cutoff and nodes were grouped manually according to association with specific GO categories flagged by STRING annotations. Interaction networks of Piwi, HP1a, Egg, Wde and Vasa were retrieved from FlyBase and nodes present in the diGly SUMO dataset were custom colored in CytoScape^48^.

### Detection of protein SUMOylation by Immunoprecipitation (IP) and Western blotting

All samples were prepared in parallel with respective controls. Hand dissected ovaries from 100-150 flies (depending on the expressed protein) of appropriate genotypes were lysed in RIPA buffer supplemented with Complete Protease Inhibitor cocktail (Roche, 11836170001) and 20 mM NEM. Debris were removed by centrifugation at 20,000 rpm for 10 minutes at 4°C. Cleared lysates were incubated with GFP-Trap magnetic agarose beads (ChromoTek, gtma-20) for 1-2 hrs with end-to-end rotation at 4°C. Beads were washed 5 times with “harsh” wash buffer (20 mM Tris pH 7.4, 0.5 M NaCl, 1% NP40 (Igepal), 0.5% Sodium deoxycholate, 1% SDS). Finally, samples were transferred to fresh tubes and boiled in LDS sample buffer (Invitrogen, NP0007) for 5 minutes at 95°C. For optimal separation of high molecular weight SUMOylated species, IP samples were analyzed on 3–8% tris-glycine gels (Invitrogen). Additionally, input samples were separately analyzed at higher percent gels to capture low molecular weight free SUMO. After electrophoresis, proteins were transferred to 0.45μm nitrocellulose membrane (BioRad) and transfer was verified by Ponceau S stain. Membranes were then blocked with 5% nonfat milk in PBS-T(0.1% Tween in PBS) for >30 min at room temperature (RT), followed by incubation with primary antibodies for 2 hrs at RT or overnight at 4°C. The following antibodies were used for detection: HRP-conjugated anti-Flag antibody (Sigma, A8592), rabbit anti-GFP (Abcam, ab290), mouse anti-Tubulin (Sigma, T5168), mouse anti-Piwi (Santa Cruz, sc-390946), HRP-conjugated anti-mouse IgG (Cell Signalling, 7076), HRP-conjugated anti-rabbit IgG (Cell Signalling, 7074), IRDye® 800CW anti-rabbit IgG (Licor). For SUMOylation analysis in S2 cells, plasmids encoding GFP-tagged Piwi, Mael, Spn-E and Panx and 3xFlag3xHA-tagged SUMO under the control of ubiquitin and actin promoter, respectively, were co-transfected using TransIT-Insect Transfection reagent (Mirus, MIR6105). Cells were harvested 48 hrs post-transfection and samples were processed and analyzed following the procedure described above.

### RT-qPCR

RNA was extracted from 20-25 pairs of ovaries from appropriate genotypes using Trizol (Invitrogen, 15596018) according to the manufacturer instructions. Purified RNA was treated with DNAse and subjected to reverse transcription with the SuperScript IV with ezDNAse kit (Invitrogen, 11766050) according to the manufacturer instructions. qPCR was performed using SybrGreen and the following primers: Su(var)2-10: CCAGCACAGGACGAACAGCCC and CGTGGAACTGGCGACGGCTT, Piwi: CTGCTGATCTCCAAAAATAGGG and TCGCGTATAACTGCTCATGG; Rp49 (internal control): CCGCTTCAAGGGACAGTATCTG and ATCTCGCCGCAGTAAACGC. Relative expression was calculated with respect to the Rp49 expression using the delta Ct method.

### Protein localization imaging

Ovaries were dissected in cold PBS and fixed in 4% formaldehyde in PBS-T (0.1% Tween-20 in PBS) with end-to-end rotation for 20 minutes at room temperature, following by 3 washes with PBS-T for >10 minutes each. For Vasa detection, fixed ovaries were incubated with blocking solution (0.3% Triton-X, 0.1% Tween-20, 3% Bovine serum albumin (BSA)) for 1 hour with end-to-end rotation, followed by an overnight incubation with 1:10 dilution of anti-vasa antibody (DSHB anti-vasa supernatant, AB_760351) at 4°C with rotation. Next, ovaries were washed 3 times with washing solution 0.3% Triton-X, 0.1% Tween-20 in PBS) for 10 minutes and incubated with 1:400 diluted AlexaFlour 594 secondary antibody (Invitrogen A-11007) in blocking solution for 2 hours at RT in the dark. Finally, ovaries were washed with washing solution for 10 minutes 3 times, 1 μg/mL DAPI solution for 5 minutes, and a final rinse of 5 minutes. For proteins GFP-Aub, fixed ovaries were only stained with DAPI. Samples were mounted on glass slides in Vectashield Antifade Mounting Medium (Vectorlabs, H-1200-10) and analyzed on an Upright Zeiss LSM 880 Confocal Microscope at the UCR Microscopy and Imaging Facility. Final images were processed and assembled with Fiji^49^. Shown are representative images of 10 or more analyzed ovaries.

### Fertility test

10 2-day old females of appropriate genotype and 5 wild type males were maintained at standard fly media for 3 and 5 days, and the numbers of viable adult offspring from each vial was manually counted. Results are presented as viable offspring per day. Additionally, vials where females expressing shRNAs against Piwi and Su(var)2-10 from different crosses were maintained prior dissection were inspected for viable progeny to verify expected sterility phenotypes.

### Data availability statement

The mass spectrometry raw data are deposited to the ProteomeXchance Consortium (https://www.ebi.ac.uk/pride/) via the PRIDE repository with the dataset identifier PXD037421. Reviewer account details: Username: xxx Password: xxx.

## Supporting information

Supplemental Material Archive

## Acknowledgements

We thank former Caltech Protein Exploration Laboratory (PEL) members Dr. Michael Sweredoski and Dr. Annie Moradian for their advice on diGly proteomics, and Corinne Karalun (former laboratory assistant in MN laboratory, UCR), Hannah Ryon (former rotation student in KFT laboratory, Caltech), and Matea Ibrahim (undergraduate students in UC Riverside) for assistance with Western Blotting, fly dissections and genotyping. This work was supported by grants from the NIH (K99/R00 HD099316) to MN; the NIH (R01 GM097363) and the Howard Hughes Medical Institute Faculty Scholar Award to AAA, and the NIH (R01 GM110217) and Ellison Medical Foundation Awards to KFT.

## Authors contributions

MN, KFT, and AAA conceptualized the proteomics study. MN designed and performed experiments, data curation, and formal analysis, except LC-MS/MS runs and raw data processing which were performed at the Caltech PEL facility by BL. HH performed GFP-Aub localization experiments. MN prepared figures and drafted the manuscript. MN and AAA edited the manuscript.

