## Supplemental Material Archive for "Pervasive SUMOylation of heterochromatin and piRNA pathway proteins": Ninova_et_al_suppl_info.pdf

#### Supplementary Information

##### Supplementary Figure Legends

**Figure S1.** Related to Figure 2B. Amino acid frequencies flanking predicted K-ε-GG for sites with high and low SUMO-TR:control intensity ratios in each replicate.

**Figure S2.** Related to Figure 2C. Relationship of diGly site intensity and reproducibility of detection. Boxplots show the distributions of intensities for all diGly sites detected in each replicate, divided into categories depending on their detectability in other replicates.

**Figure S3.** Related to Figure 2F. Localization of SUMOylation sites to IUPred2A-perdicteed protein IDRs. Boxplots show the median IDR score for 10-aminoacid region flanking the central diGly modified lysine for bona fide SUMO sites, compared to randomly chosen lysine in the same protein and to background diGly sites.

**Figure S4.** Related to Figure 4. Confocal images of ovarioles from control and SUMO knockdown via MT-gal4 driven shRNA ovaries stained for DAPI. The arrowhead indicates collapsed nuclei in mid-stage egg chambers in SUMO knockdown ovaries.

**Figure S5.** Related to Figure 5. Western blot analysis (WB) protein SUMOylation in *Drosophila* S2 cells. Indicated GFP-tagged proteins were overexpressed in S2 cells alongside 3xFlag3xHA tagged SUMO, and cell lysates were subjected to GFP immunoprecipitation followed by WB detection first with anti-Flag antibody, and after stripping, anti-GFP antibody. Asterisks indicate the full length unmodified GFP-tagged protein form (red) or putative SUMO-modified form (green). Arrows indicate putative degradation products of unmodified (red) and SUMO-modified (green) GFP-tagged protein. Images are representative from two independent biological replicates.

##### Supplementary Tables

**Table S1.** MaxQuant GlyGly site data table (available with PRIDE repository with the dataset identifier PXD037421)

**Table S2.** GO analysis data

### Supplementary Figure 1

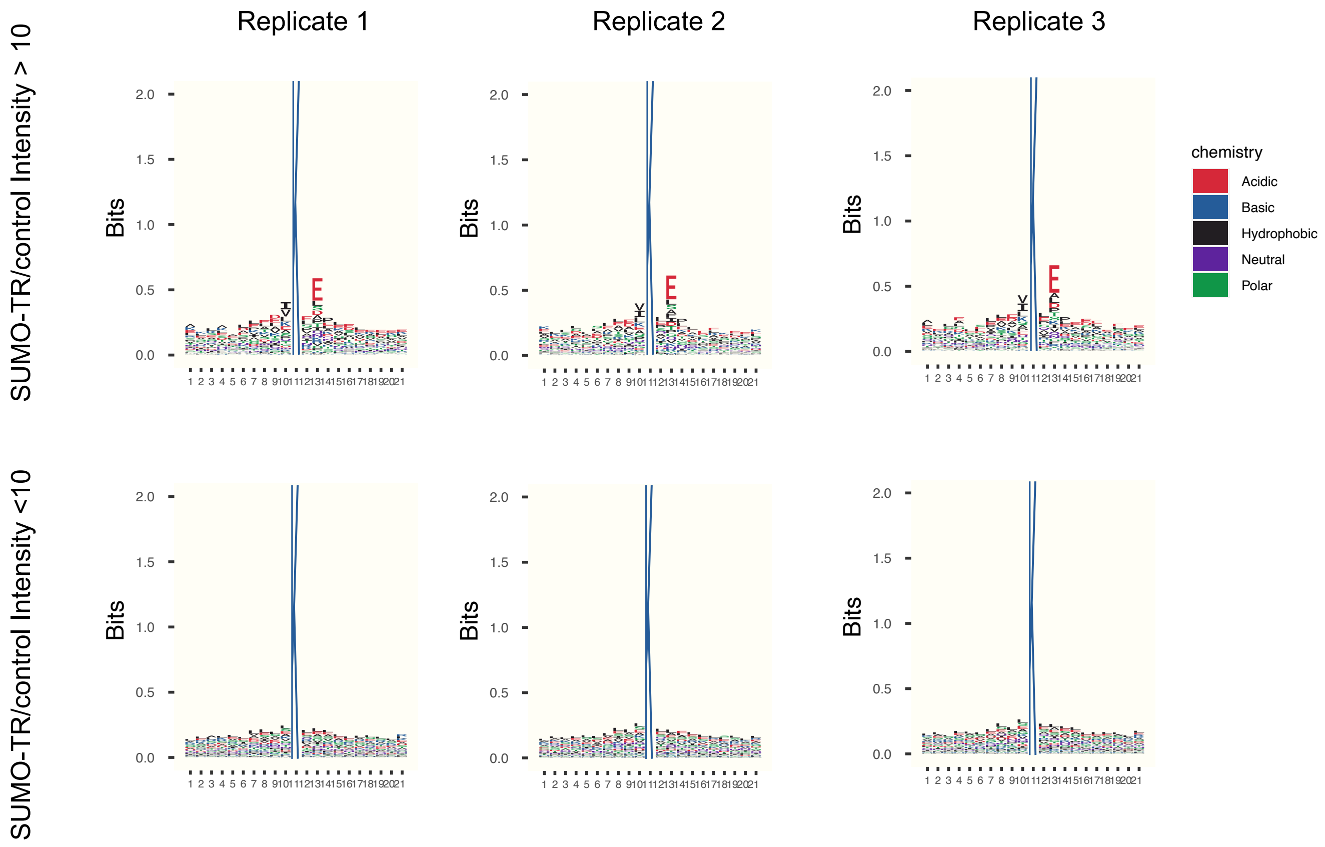

### Supplementary Figure 2

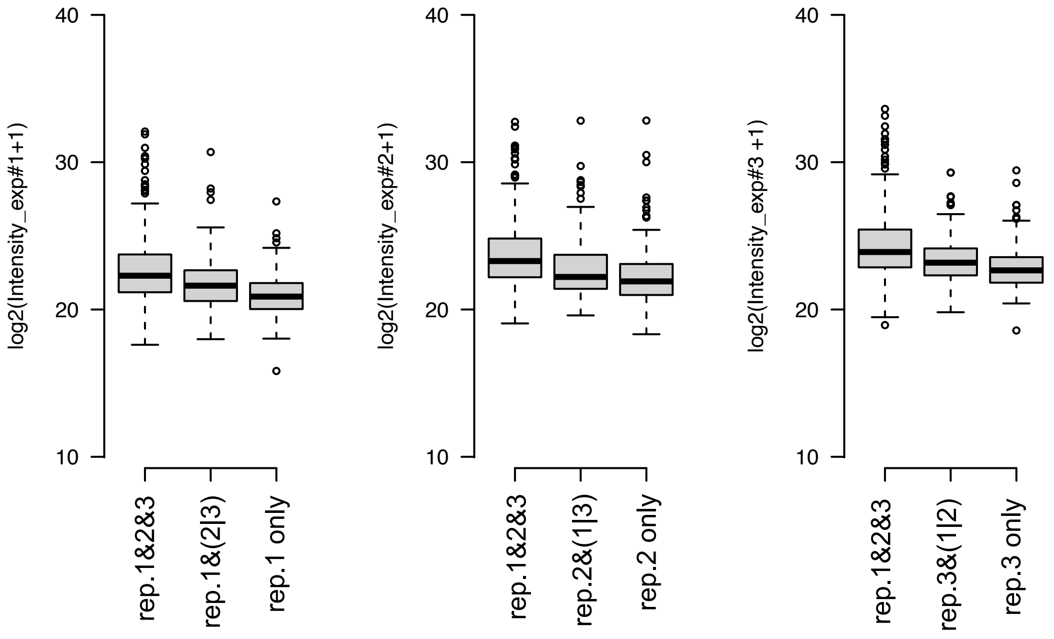

### Supplementary Figure 3

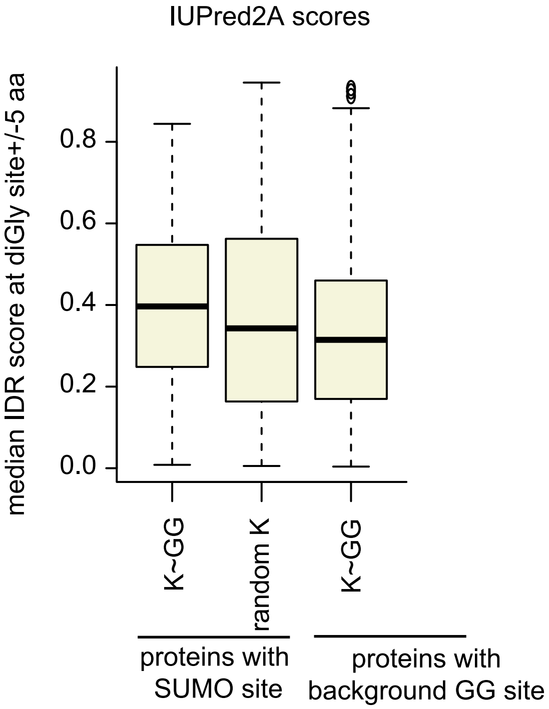

### Supplementary Figure 4

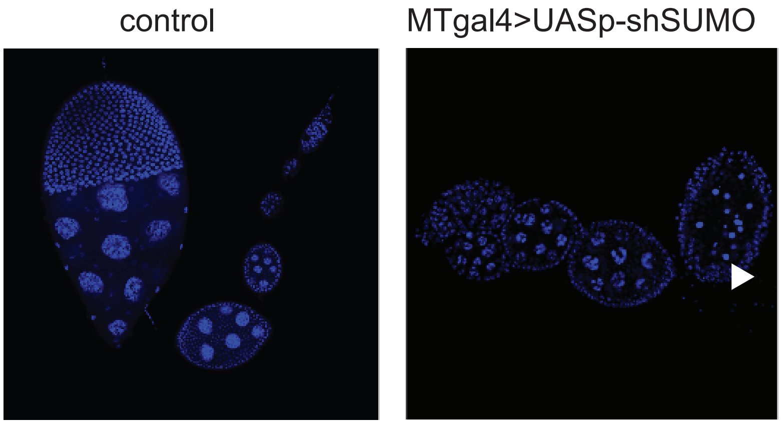

### Supplementary Figure 5

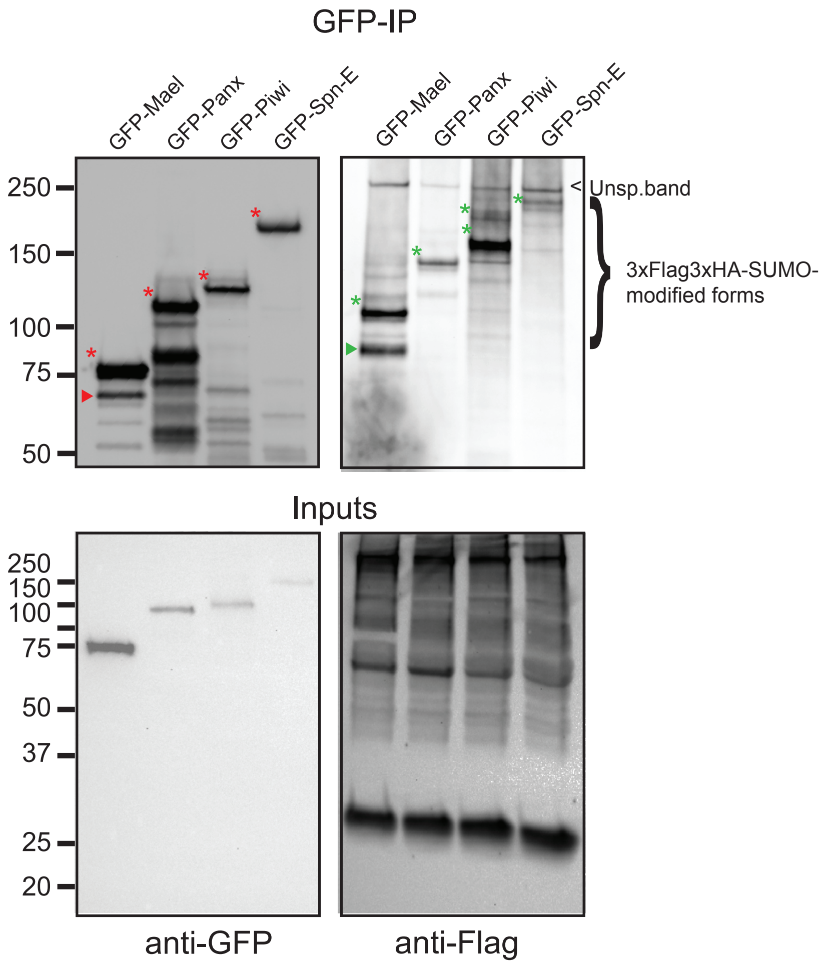
